# One-step N-terminomics based on isolation of protein N-terminal peptides from LysargiNase digests by tip-based strong cation exchange chromatography

**DOI:** 10.1101/2024.02.06.579163

**Authors:** Kazuya Morikawa, Hiroshi Nishida, Koshi Imami, Yasushi Ishihama

## Abstract

We have developed a one-step isolation method for protein N-terminal peptides from LysargiNase digests by pipette tip-based strong cation exchange (SCX) chromatography. This CHAMP-N (CHromatographic AMplification of Protein N-terminal peptides) method using disposable and parallel-processable SCX tips instead of conventional HPLC SCX columns facilitates simple, sensitive and high-throughput N-terminomic profiling without sacrificing the high identification numbers and selectivity achieved by the HPLC-based method. By applying the CHAMP-N method to HEK293T cells, we identified novel cleavage sites for signal and transit peptides, and non-canonical translation initiation sites. Finally, for proteome-wide terminomics, we present a simple and comprehensive N-and C-terminomics platform employing three different tip-based approaches, including CHAMP-N, in which protease digestion and one-step isolation by tip LC are commonly used to achieve complementary terminome coverages.

## Introduction

Proteins encoded by a single gene may have various sequences and modifications generated by multiple mechanisms, including alternative splicing, mutation, translational control, proteolysis and co- or post-translational modifications. These diverse protein isoforms are called proteoforms (1–3). Each proteoform generally has a distinct function, localization, and stability (4). For example, it has been reported that proteoforms derived from non-canonical translation initiation sites affect tumorigenesis (5–7), and that proteoforms cleaved at different sites of the amyloid precursor protein are involved in the development of Alzheimer’s disease (8). Each of these proteoforms resulting from different translation initiation sites or proteolytic cleavages has a different protein terminal sequence. Moreover, localization and stability are affected by signal or transit peptide sequences and N- or C-degrons (9) present at the protein termini. Thus, comprehensive analysis of protein termini, or terminomics, is essential for understanding the functions of diverse proteoforms.

In terminomics, protein terminal peptides are isolated from other digested non-terminal peptides and subjected to liquid chromatography/tandem mass spectrometry (LC/MS/MS). Many methods to isolate protein N- and C-terminal peptides have been developed. COFRADIC (combined fractional diagonal chromatography) (10) and TAILS (terminal amine isotopic labeling of substrates) (11) are widely known N-terminal peptide isolation methods, and other methods for N-terminal peptide isolation have also been reported (12). However, all of these methods require complex and time-consuming procedures involving chemical modifications to block amines, and the efficiency and selectivity of these chemical modifications are still problematic. Instead, we have recently developed simple isolation methods for protein N- and C-terminal peptides that consist only of enzymatic digestion of proteins followed by chromatographic separation; we call these methods CHromatographic AMplification of Protein N- and C-terminal peptides (CHAMP-N (13) and CHAMP-C (14)). With these methods, terminal peptides can be isolated with high selectivity in a single step, avoiding the need for the complex steps required in the conventional methods. In the CHAMP-C method, C-terminal peptides can be easily isolated from V8 protease digests using pipette-tip columns packed with CeO_2_ particles. On the other hand, the CHAMP-N method, which uses HPLC-based strong cation exchange (SCX) chromatography for LysargiNase digests, can be fully automated, but this requires dedicated equipment and has low-throughput because it analyzes one sample at a time in series.

In this study, we developed a simple, high-throughput protein N-terminal peptide isolation method from LysargiNase digests using SCX pipette-tips instead of HPLC columns, and applied it to the identification of novel post-translational cleavage sites (signal peptide and transit peptide cleavage sites) and non-canonical translation initiation sites. We also established a comprehensive terminomics platform by combining the developed LysargiNase/SCX tip-based CHAMP-N method with our previously reported V8 protease/CeO_2_ tip-based CHAMP-C method (14) and the trypsin/SCX tip-based CHAMP-NC method (15–19).

### Experimental procedures

#### Materials

LysargiNase was purchased from Merck Millipore (Burlington, VT). Empore cation-SR disks and SDB-XC Empore disks were purchased from GL Sciences (Tokyo, Japan). UltraPure Tris Buffer and fetal bovine serum were purchased from Thermo Fisher Scientific (Waltham, MA). Protease inhibitors were purchased from Sigma-Aldrich (St. Louis, MO). TPCK-treated sequencing-grade modified trypsin was obtained from Promega (Madison, WI). Polyethylene frit was purchased from Agilent Technologies (Santa Clara, CA). Water was purified using a Millipore Milli-Q system (Bedford, MA). All other chemicals were purchased from FUJIFILM Wako (Osaka, Japan), unless otherwise specified.

#### Cell Culture, Protein Extraction, and Lysarginase Digestion

HEK293T cells from the RIKEN BRC cell bank (Ibaraki, Japan) were cultured in Dulbecco’s modified Eagle’s medium with 10% fetal bovine serum at 37 °C under 5% CO_2_. Proteins were extracted with a phase-transfer surfactant (PTS protocol), as described previously (20), except that LysargiNase was used instead of trypsin/LysC. Briefly, HEK293T cells were suspended in PTS lysis buffer consisting of protease inhibitors, 12 mM sodium deoxycholate (SDC), 12 mM sodium *N*-lauroylsarcosinate (SLS) in 100 mM Tris−HCl buffer (pH 9). The cell lysate was diluted tenfold with 10 mM CaCl_2_ solution and incubated overnight with LysargiNase [enzyme/substrate ratio = 1:50 or 1:100 (w/w)] at 37 °C. After digestion, an equal volume of ethyl acetate was added, and the solution was acidified with trifluoroacetic acid (TFA). Then, the peptides were desalted with SDB-StageTips (21, 22).

#### Strong Cation Exchange Chromatography

SCX chromatography was performed using disposable pipette tip-based SCX-StageTips (22), prepared as follows: double SCX membrane from Empore cation-SR disks was stamped out using a blunt-end 16-gauge syringe needle and placed into a P200 pipette tip. Mobile phases were prepared with aqueous 30% acetonitrile in buffers with various acid or salt solutions (Table S1). Prior to use, SCX-StageTips were activated with aqueous 30% acetonitrile in 1 M NaCl, and equilibrated with 300 μL of the mobile phase. Ten microgram aliquots of desalted peptide samples were dried in a SpeedVac concentrator (Thermo Fisher Scientific), reconstituted with the mobile phase, and each sample was loaded onto an SCX-StageTip. The eluted fractions were collected. After SpeedVac concentration, each sample was reconstituted in 4% acetonitrile with 0.5% TFA and injected onto a nanoLC/MS/MS system equipped with an Orbitrap Fusion Lumos mass spectrometer (Thermo Fisher Scientific) or Orbitrap Exploris 480 mass spectrometer (Thermo Fisher Scientific), as described below. To compare HPLC-based and tip-based CHAMP-N, peptide isolation experiments were performed in triplicate using 80 μg x 3 of LysargiNase-digested HEK293T peptides. A quarter of the isolated peptides were injected into the nanoLC/MS/MS system using Orbitrap Fusion Lumos and measurements were performed in triplicate.

#### Isolation of C-terminal and N, C-terminal peptides by CHAMP-C and CHAMP-NC

C-Terminal peptide isolation by CHAMP-C (14) and N, C-terminal peptide isolation by CHAMP-NC (15, 19) were performed as described previously. In brief, C-terminal peptide isolation was performed using CeO_2_ tips, which were loaded with methanol-suspended CeO_2_ particles and centrifuged on a P200 pipette tip with a polyethylene frit. Ten microgram aliquots of the V8 protease-digested peptide samples were reconstituted with 40% acetonitrile containing 50 mM 5-aminovaleric acid-HCl and 1 M NaCl, and each sample was loaded onto a CeO_2_ tip. N, C-Terminal peptide isolation was performed using SCX-StageTips. Ten microgram aliquots of the trypsin/Lys-C-digested peptide samples were reconstituted with 30% acetonitrile containing 0.125 % TFA, and each sample was loaded onto an SCX-StageTip. In both methods, the eluted fractions were collected.

#### NanoLC/MS/MS Analysis

The LC/MS/MS analyses were performed on an HTC-PAL autosampler (CTC Analytics, Zwingen, Switzerland) with an Ultimate 3000 pump (Thermo Fisher Scientific), combined with an Orbitrap Fusion Lumos mass spectrometer or an Ultimate 3000 liquid chromatograph combined with an Orbitrap Exploris 480 mass spectrometer (Thermo Fisher Scientific). The LC mobile phases consisted of solvent A (0.5 % acetic acid) and solvent B (0.5 % acetic acid and 80 % acetonitrile).

For the Orbitrap Fusion Lumos system, the peptides were separated on self-pulled needle columns (150 mm, 100 μm ID) packed with Reprosil-Pur 120 C18-AQ 3 μm (Dr. Maisch, Ammerbuch, Germany) (23), and separated by a linear gradient: 5-10 % B in 5 min, 10-40 % B in 60 min, and 40-99 % B in 5 min, followed by 99 % B for 10 min. The flow rate was 500 nL/min. The electrospray voltage was set to 2.4 kV in the positive mode. The mass range of the survey scan was from 300 to 1,500 *m/z* with resolution of 120,000, standard automatic gain control (AGC) target, and maximum injection time of 50 ms. The first mass of MS/MS scan was set to 110 *m/z* with a resolution of 15,000, 100% normalized AGC target, and maximum injection time of 50 ms. The fragmentation was performed by higher energy collisional dissociation with a normalized collision energy of 30 %. The dynamic exclusion time was set to 20 s.

For the Orbitrap Exploris 480 system, the peptides were separated on self-pulled needle columns (250 mm, 100 μm ID) packed with Reprosil-Pur 120 C18-AQ 1.9 μm (Dr. Maisch, Ammerbuch, Germany) at 50 °C in a column oven (Sonation), and separated by a linear gradient: 5 % B in 8.3 min, 5-19 % B in 92.2 min, 19-29 % B in 34.5 min, 29-40 % B in 15 min, and 40-99 % B in 0.1 min, followed by 99 % B for 4.9 min. The flow rate was 400 nL/min. The electrospray voltage was set to 2.4 kV in the positive mode. The mass spectrometric analysis was carried out with the FAIMS Pro interface. The FAIMS mode was set to a standard resolution, and the total carrier gas flow was 4.0 L/min. The compensation voltage (CV) was set to −40, −60, and − 80, and the cycle time of each CV experiment was set to 1 s. The mass range of the survey scan was from 300 to 1,500 *m/z* with resolution of 60,000, 300% normalized AGC target, and auto maximum injection time. The first mass of MS/MS scan was set to 120 *m/z* with a resolution of 30,000, standard AGC target, and auto maximum injection time. The fragmentation was performed by higher energy collisional dissociation with a normalized collision energy of 30 %. The dynamic exclusion time was set to 20 s.

#### Data Analysis

The raw MS data files acquired with FAIMS were converted into an MzXML format using FAIMS MzXML Generator (https://github.com/coongroup/FAIMS-MzXML-Generator) (24). The obtained mzxml files as well as the raw MS data files acquired using the Orbitrap Fusion Lumos mass spectrometer were searched by MaxQuant (ver. 1.6.17.0) (25), and the database search was performed with Andromeda (26) against the SwissProt database (ver. 2020-10, 42,372 human protein entries). For most of the CHAMP-N experiments, search parameters included two missed cleavage sites for LysargiNase-digested peptides, while LysargiNase-digestion with N-terminal free semi-specificity was set for the searching of Neo-N-termini by CHAMP-N. Two missed cleavage sites for trypsin-digested peptides were set for the CHAMP-NC experiments, and two missed cleavage sites for V8 protease-digested peptides were set for the CHAMP-C experiments. Protein N-terminal acetylation and methionine oxidation were set as variable modifications, and cysteine carbamidomethylation was set as a fixed modification. For the searching of Neo-N-termini by CHAMP-N, peptide N-terminal acetylation was set instead of protein N-terminal acetylation. The peptide mass tolerance was 4.5 ppm, and the MS/MS tolerance was 20 ppm. The false discovery rate was set to 0.01 at the peptide-spectrum match level and protein level.

### Experimental Design and Statistical Rationale

All experiments were performed using aliquots of the same HEK293T digest to minimize confounders involving steps prior to isolation of the terminal peptides. Isolation of terminal peptides was prepared with n = 3, and in most cases a single LC/MS/MS measurement was performed, whereas for comparison with the HPLC-based method the LC/MS/MS measurements were performed in triplicate for each sample. No statistical analysis was performed in this study.

## RESULTS AND DISCUSSION

### Optimization of Isolation Conditions for N-Terminal Peptides Using Pipette-tip SCX Columns

This study aimed to establish the tip-based CHAMP-N method, in which N-terminal peptides are isolated with high selectivity using a pipette-tip microcolumn (Figure S1A) instead of the conventional HPLC-based method (13). In this method, proteins are initially digested with LysargiNase, a digestive enzyme that cleaves the N-terminal side of Lys and Arg. Peptides digested with LysargiNase have 0 charge on acetylated N-terminal peptides and +1 charge on unmodified N-terminal peptides, whereas non-N-terminal peptides have at least +2 charge under acidic conditions. Based on the charge/orientation retention model (17), non-N-terminal peptides with two positive charges in close proximity are strongly retained in the SCX. Thus, due to the relatively weak retention, acetylated N-terminal peptides with 0 charge, as well as unmodified N-terminal peptides with +1 charge, and N-terminal peptides with +2 charge (e.g., His-containing peptides) with the charges at distant positions can be simultaneously isolated (Figure S1B). We have previously reported the CHAMP-N method using a pipette-tip microcolumn (15, 19). However, we found that N-terminal peptides with +2 charge could not be isolated under the isolation conditions with formic acid (FA) used in that work (Figure S2A, B). Even when the isocratic elution volume was increased, N-terminal peptides with +2 charge were hardly isolated (Figure S2C, D). In this study, we first investigated suitable elution conditions to elute N-terminal peptides with high selectivity and without loss of +2 charge N-terminal peptides, using HEK293T cell-derived proteins.

Salt- and pH-based elution are two major modes for peptide separation by SCX chromatography (27, 28). To isolate protein N-terminal peptides by SCX-HPLC (13), we employed salt-based elution under acidic conditions. Previously, Adachi *et al.* reported the superiority of strong acid-based elution compared to salt-based elution in an SCX-StageTip-based fractionation (29). Therefore, we compared salt-based elution with acid-based elution. For the salt-based elution, we used the same eluents as those used in the HPLC-based method (13). For the acid-based elution, an eluent containing 0.2 % TFA, which is a stronger acid than FA used in the above experiment, was used. Isocratic elution was conducted and 100 μL of the eluent was collected for each fraction. Six fractions were collected and measured by LC/MS/MS. We found that the peptides eluted in the order of the charge/orientation retention model under both conditions, and protein N-terminal peptides were successfully isolated (Figure 1A). Although the elution profiles were similar under both conditions, more N-terminal peptides were identified by acid-based elution than by salt-based elution (Figure 1B, C). This result is consistent with the previous report in which acid-based elution showed higher separation efficiency than salt-based elution in peptide fractionation (29).

**Figure 1.**
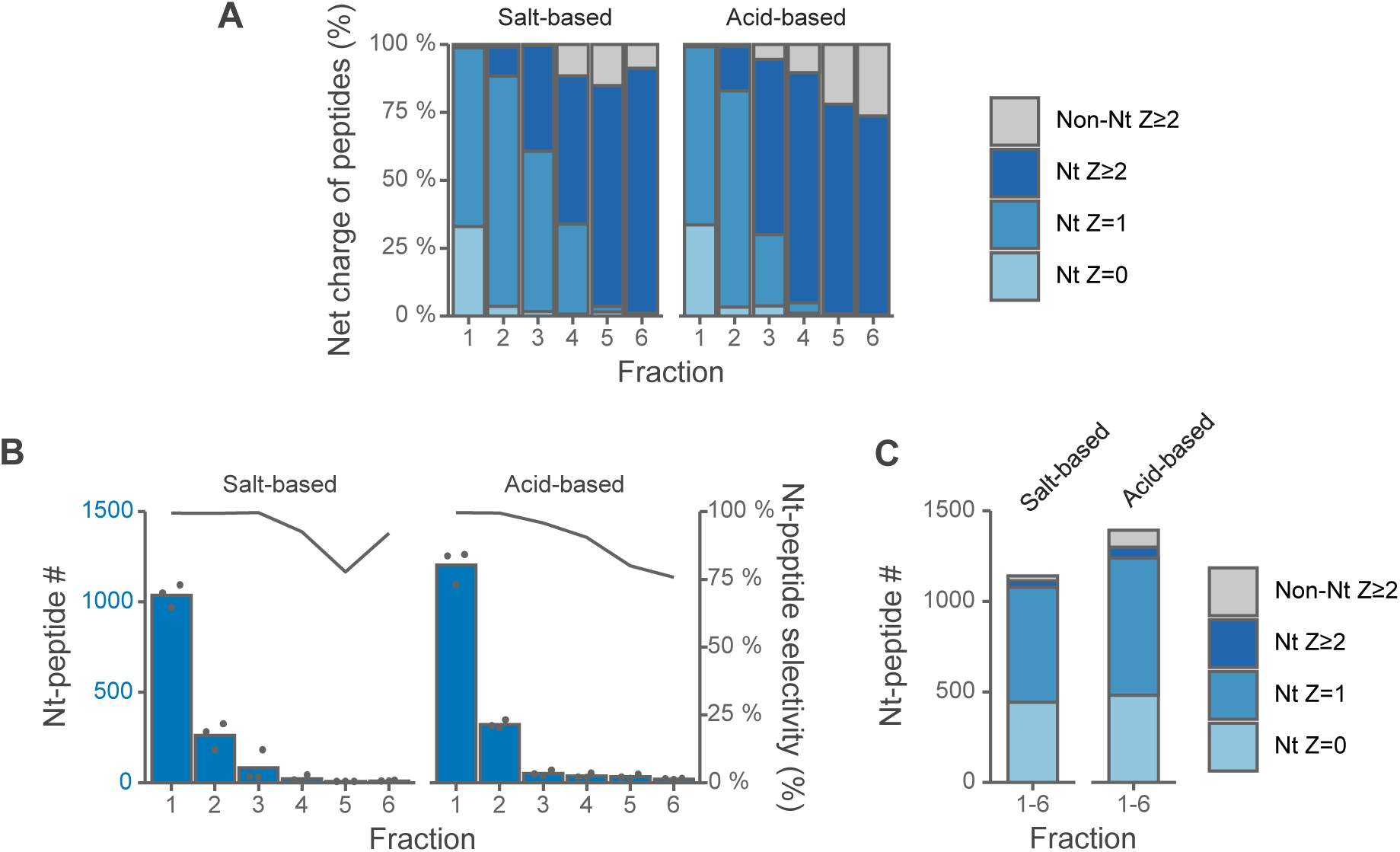
Comparison of salt-based and acid-based isocratic elution of protein N-terminal peptides. SCX tip-based separation of LysargiNase-digested HEK293T peptides under isocratic conditions with salt (10 mM KCl, 7.5 mM potassium phosphate buffer at pH 2.2 and 30% ACN) or acid (0.2% TFA and 30% ACN). Z is the charge number at acidic pH, which is based on the number of basic residues per peptide, such as unmodified N-terminus, Lys, Arg and His. Each eluted fraction was 100 µL. The Orbitrap Fusion Lumos system was used. (A) Distributions of Z values of identified peptides. (B) Numbers of identified N-terminal peptides. N-Terminal peptide selectivity (%) was calculated as the sum of the signal intensity of all identified N-terminal peptides divided by that of all identified peptides. (C) Numbers of peptides identified in any of the six fractions.

When the TFA concentration was increased from 0.2 % to 0.5 %, N-terminal peptides were isolated at a smaller elution volume. After fraction 3, the selectivity decreased sharply (Figure 2A, B), and 96.4 % of all N-terminal peptides identified, or 1794 N-terminal peptides, were identified from fractions 1 or 2 (Figure 2C). These results indicate that N-terminal peptides can be isolated selectively without loss by collecting fractions 1 and 2 (100 μL in total) under the 0.5 % TFA condition.

**Figure 2.**
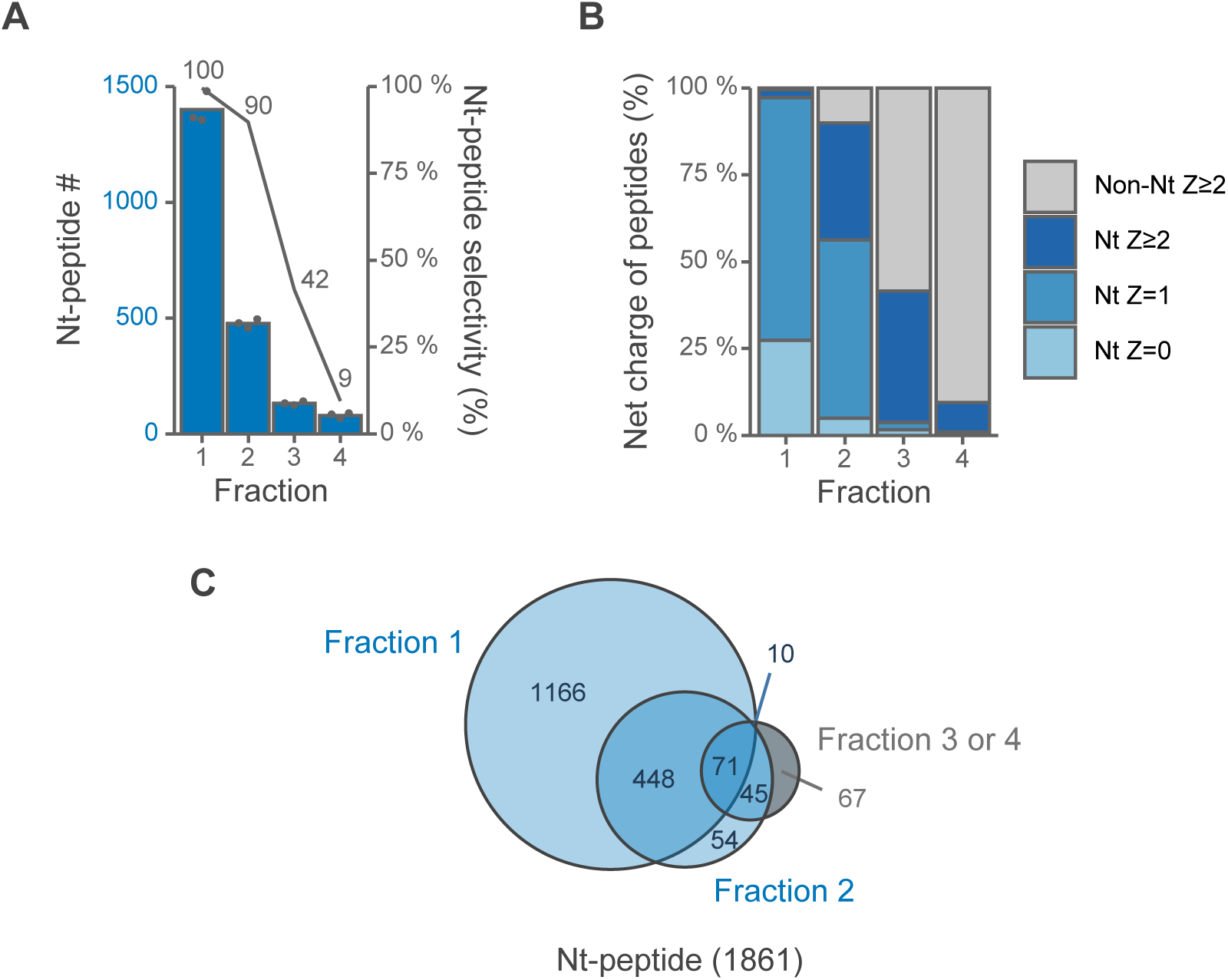
Optimization of the elution volume for SCX tip-based separation of LysargiNase-digested HEK293T peptides under acid-based isocratic condition (0.5% TFA and 30% ACN). Z is the charge number at acidic pH, which based on the number of basic residues per peptide, such as unmodified N-terminus, Lys, Arg and His. Each eluted fraction was 50 µL. The Orbitrap Fusion Lumos system was used. (A) Numbers of identified protein N-terminal peptides. N-Terminal peptide selectivity (%) was calculated as the sum of the signal intensity of all identified N-terminal peptides divided by that of all identified peptides. (B) Distributions of Z values of identified peptides. (C) Overlaps of identified N-terminal peptides in each fraction.

Based on the N-terminal peptide isolation conditions with salt- and acid-based elution described above, we compared the identification number and selectivity of N-terminal peptides in a single LC/MS/MS run without pre-fractionation. We found that more N-terminal peptides were identified by acid-based elution than by salt-based elution, as was the case with fractionation (Figure S3A). N-Terminal peptides could be isolated with a high selectivity of more than 95 % under all conditions (Figure S3A, B). The optimal conditions were those of acid-based elution with 0.5 % TFA, which allowed identification of a larger number of N-terminal peptides with a smaller elution volume. Next, we compared the conditions using 0.5 % TFA with those using FA previously reported (15, 19). Compared to the conventional method with 2.5 % FA, the present method with 0.5 % TFA identified more N-terminal peptides (Figure S3C). In addition, N-terminal peptides with +2 charge could be isolated by this method (Figure S3D), whereas they could hardly be isolated by our previous method. In subsequent experiments, therefore, acid-based elution with 0.5% TFA was used.

### Comparison of Conventional HPLC-based Method and the Tip-based Method

Next, we compared the performance of the new tip-based method with that of the previously reported HPLC-based method (13). The results of the HPLC-based method for comparison were obtained by reanalyzing the previously published data (PXD010551) using the same database search parameters as employed for the tip-based method. The HPLC-based method identified 1875 N-terminal peptides on average, whereas the tip-based method identified 1985 N-terminal peptides on average in a single LC/MS/MS run (Figure 3A). The tip-based method showed relatively lower variability among replicates, which may be attributed to the fact that, unlike the HPLC-based method, multiple tips can be used for parallel processing. On the other hand, N-terminal peptides with two positive charges were detected at a higher rate by the HPLC-based method (Figure 3B). These results suggest that it is difficult to separate non-N-terminal peptides with two positive charges located close to each other from N-terminal peptides with two positive charges located far apart, and that the HPLC column is superior to the tip-based method in this respect. Thus, the tip-based method is inferior to the HPLC-based method in a few aspects. However, the tip-based method was able to isolate N-terminal peptides with a high selectivity of more than 92%, and considering the advantages of easy operation and high throughput, we believe it represents a practical and simple protocol.

**Figure 3.**
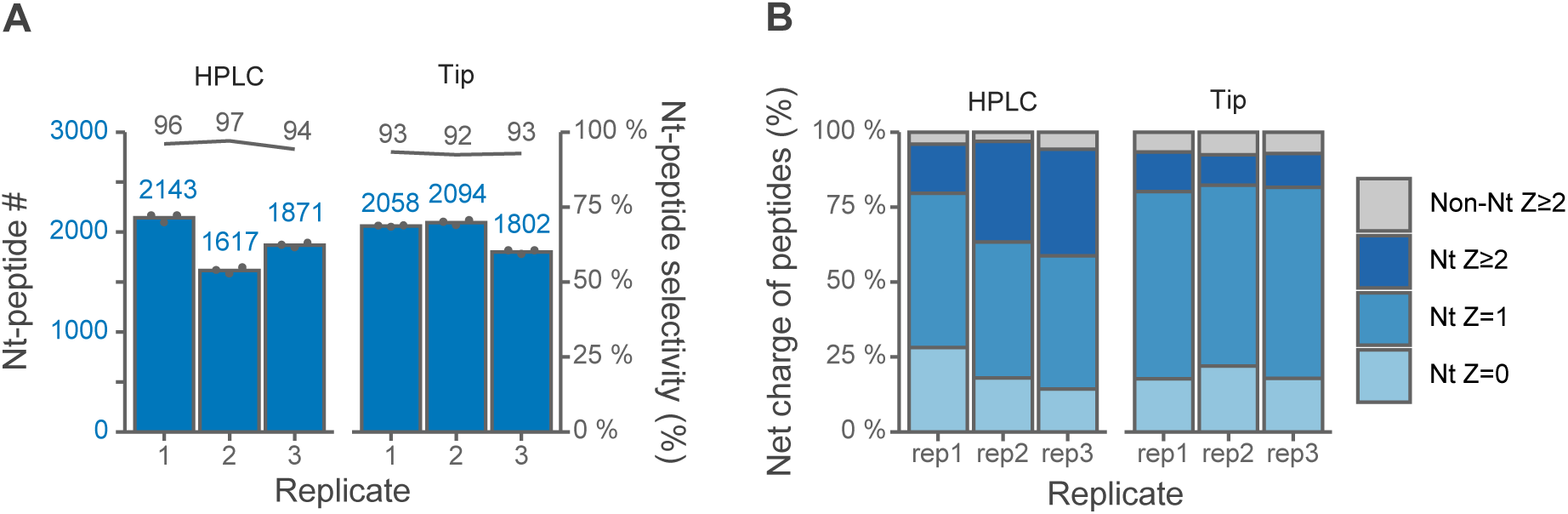
Comparison of HPLC-based and tip-based isolation of protein N-terminal peptides. SCX HPLC or SCX tip separation of LysargiNase-digested HEK293T peptides. Z is the charge number at acidic pH, which based on the number of basic residues per peptide, such as unmodified N-terminus, Lys, Arg and His. The HPLC results were obtained by reanalysis of data in our previous report using Orbitrap Fusion Lumos (JPST000422). For each replicate, three LC/MS/MS runs were performed. The Orbitrap Fusion Lumos system was used. (A) Numbers of identified N-terminal peptides. N-Terminal peptide selectivity (%) was calculated as the sum of the signal intensity of all identified N-terminal peptides divided by that of all identified peptides. (B) Distributions of Z values of identified peptides.

### Identification of Proteolytic Cleavage Sites and Translation Initiation Sites

We applied this tip-based CHAMP-N method to the detection of neo-N-termini derived from proteolysis or non-canonical translation initiation. Using “semi-specific search free N-terminus” in the MaxQuant search, peptides whose N-termini do not match the specificity of the enzyme, neo-N-terminal peptides, can be additionally identified. This means that it is possible to detect neo-N-termini resulting from downstream shifts of the translation initiation site or endogenous proteolysis, which differ from the native N-terminal sequence in the database.

As a result, over 900 neo-N-terminal peptides were identified in at least one replicate (Figure S4A). Among them, 108 of the neo-N-termini were acetylated. Since most acetylation of protein N-termini occurs co-translationally (30), these neo-N-termini are suggested to be due to non-canonical translation initiation sites. Actually, 85 of the 108 were acetylated at Met or the residue adjacent to Met (Figure 4A). The removal of the initiator methionine at the protein N-terminus and the occurrence of acetylation are dependent on the residue adjacent to Met (30). Interestingly, the frequencies of the adjacent residues in the 85 acetylated neo-N-termini were similar to those in the native N-termini (Figure 4B). Non-canonical translation initiation can occur not only from unannotated methionine (AUG), but also from AUG-like codons called near-cognate codons (31–33). Among the acetylated neo-N-termini identified in this study, near-cognate initiation codons, especially CUG-initiated translation initiation site candidates, were identified in two proteins (Figure S4B). These two candidate translation initiation sites were not identified by a recently reported approach for accurately identifying translation initiation sites, including initiation at near-cognate codons (34). That approach uses the results of applying three N-terminomics methods to HEK293T samples, and our method identified sites that were not found there.

**Figure 4.**
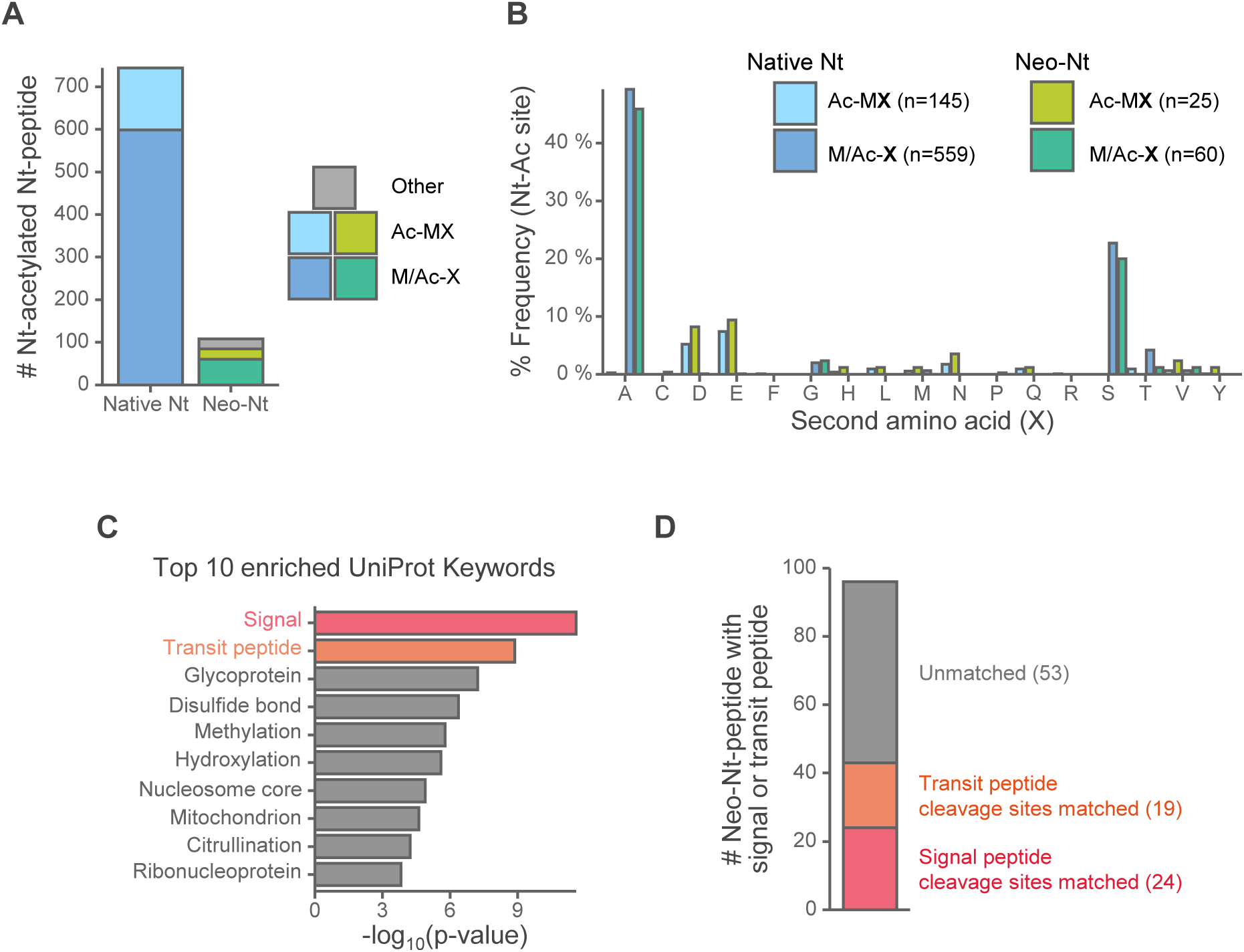
Application of tip-based CHAMP-N to identification of translation initiation sites and cleavage sites of transit peptides and signal peptides. Peptides identified in at least one of the three replicates were analyzed. The Orbitrap Exploris 480 system was used. (A) Numbers of identified Nt-acetylated Nt-peptides. The Nt-peptides are categorized into three classes. Ac-MX: The Nt-peptides acetylated at the first (initiator) methionine. M/Ac-X: The Nt-peptides acetylated at the adjacent residue to methionine. Other: The Nt-peptides without methionine around the acetylated residue. (B) The amino acid frequency at the second residue, adjacent to iMet, of acetylated protein N-termini based on the absence or presence of iMet. (C) Enrichment analysis using UniProt Keywords for the proteins with neo-Nt-peptides. (D) Numbers of transit peptide or signal peptide cleavage site matched to UniProt annotations or unmatched neo-Nt-peptides.

Another feature of the neo-N-termini was found to be that many of them are located in the N-terminal region (less than 50 residues) of the whole protein (Figure S4C). We also found that proteins containing signal and transit peptides were the most enriched with neo-N-termini (Figure 4C). These results suggest that the neo-N-termini generated by the cleavage of signal or transit peptides can be detected. Indeed, when we checked whether the lengths of the signal peptide and transit peptide sequences annotated in UniProt matched the positions of the neo-N termini identified in this study, 24 neo-N termini matched the signal peptide and 19 neo-N termini matched the transit peptide (Figure 4D). For example, COX6A1 protein localized to mitochondria has a transit peptide at residues 1 through 24, and a neo-N-terminal peptide from residue 25 was identified (Figure S5A).

Among the neo-N-termini that did not match the cleavage position annotated in UniProt, there were several neo-N-termini that deviated by a few residues from the UniProt cleavage position. Since some of the neo-N-termini annotated in UniProt lack experimental evidence, it is possible that the neo-N-termini identified in this study are the true cleavage positions. Therefore, we used TargetP 2.0 (35) as a transit peptide prediction model and SignalP 6.0 (36) as a signal peptide prediction model to search for candidate true cleavage positions. Although UniProt considers results from multiple prediction tools in the absence of experimental evidence, it is unclear whether the latest versions of SignalP and TargetP are used. The latest version, SignalP 6.0, is a signal peptide prediction model using a protein language model and has better prediction performance than SignalP 5.0 based on deep learning. Using these prediction models and experimental data from this study, we attempted to identify the true cleavage site candidates. As a result, 7 neo-N-termini matched the cleavage sites predicted by SignalP 6.0 (Table 1), and 11 neo-N-termini matched the cleavage sites predicted by TargetP 2.0 (Table 2). Most of these 18 proteins have annotated signal or transit peptide sequences in UniProt, but there is experimental evidence for only two. Therefore, for these proteins, the neo-N-terminal positions identified in this study are candidates for the true cleavage positions. For example, FKBP9 protein has a signal peptide from residue 1 to 24 annotated on UniProt, but according to the SignalP 6.0 prediction, residues 1 to 29 are the signal peptide, and we actually identified a neo-N-terminal peptide from residue 30 (Figure S5B). Of the 18 neo-N termini matched in this study, 12 are also registered in TopFIND (37), a database of protein termini, supporting the possibility that these termini are the true cleavage positions. It seems likely that more experimental information on protein termini can be obtained by this method or other terminomics methods in the future, which should lead to further improvement of the prediction model accuracy. Synergistic effects can also be expected, such as experimental validation based on the improved prediction model.

**Table 1.** Proteins with signal peptide cleavage sites consistent with SignalP 6.0 predictions.

| UniProt accession number | Gene name | SignalP 6.0 | Neo-Nt position in this study | Experimental evidence in UniProt | UniProt signal peptide | TopFIND 4.1 |
| --- | --- | --- | --- | --- | --- | --- |
| **O00115** | DNASE2 | 1-16 | 17 | No | 1-18 | - |
| **P13667** | PDIA4 | 1-24 | 25 | No | 1-20 | + |
| **P04843** | RPN1 | 1-24 | 25 | No | 1-23 | + |
| **Q14257** | RCN2 | 1-25 | 26 | No | 1-22 | + |
| **Q9Y3Q3** | TMED3 | 1-27 | 28 | No | 1-23 | + |
| **O95302** | FKBP9 | 1-29 | 30 | No | 1-24 | - |
| **P11047** | LAMC1 | 1-35 | 36 | No | 1-33 | - |

Neo-N-termini matching predicted cleavage sites by SignalP 6.0. UniProt information is current as of October 2023.

**Table 2.**
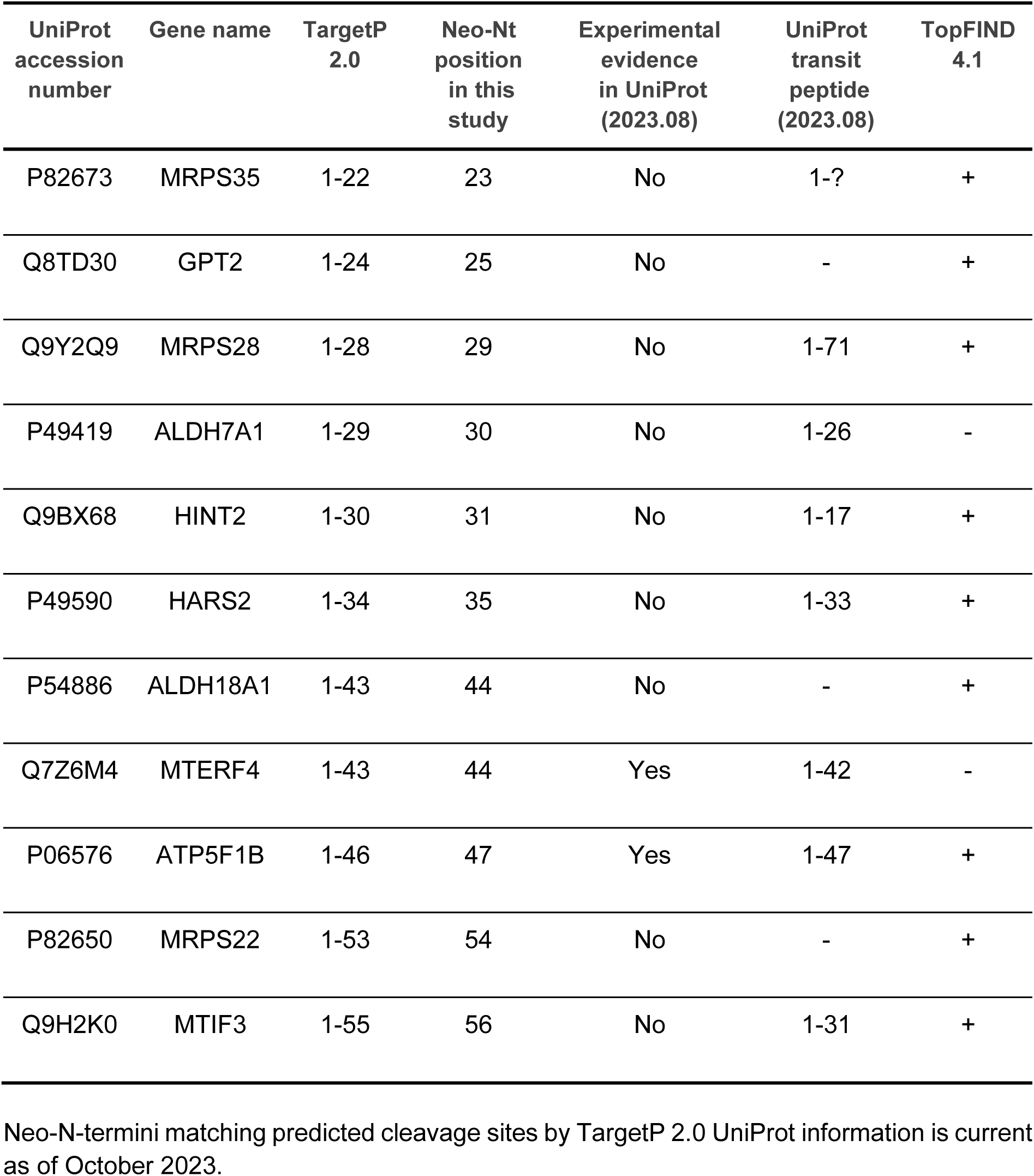
Proteins with transit peptide cleavage sites consistent with TargetP 2.0 predictions.

As mentioned above, most of the neo-N-termini identified in this study are derived from the N-terminal region of the whole protein, but the majority of the neo-N-termini are located less than 10 residues from the native N-terminus (Figure S4D). Typical signal and transit peptides are 10 to 50 residues in length, but shorter peptides are cleaved. In particular, the neo-N-terminus starting at residue 3 was the most abundant, being identified in 70 proteins. In 12 of them, the N-terminus was acetylated, suggesting that it may be excessively cleaved by methionine aminopeptidase, which removes the initiator methionine during translation. We also found that some proteins had more than 10 neo-N-terminal peptides derived from a single protein (Figure S4E). Many of these neo-N-terminal peptides were peptide ladders with one amino acid residue successively shaved off (Figure S4F), suggesting cleavage by an exopeptidase. These findings suggest that excessive cleavage by methionine aminopeptidase and cleavage by exopeptidase may occur frequently in the cell.

### Comprehensive Protein N- and C-Terminome Analysis Using Three CHAMP Methods

We recently reported the CHAMP-C method (14), which is a tip-based method for isolation of C-terminal peptides, and the CHAMP-NC method (15, 19), which is a tip-based version of a previously developed method for simultaneous isolation of N- and C-terminal peptides (Figure 5A). The CHAMP-C method consists only of protein digestion by V8 protease followed by metal oxide-based ligand-exchange chromatography (MOLEX). When digested by V8 protease, which cleaves the C-terminal side of Asp and Glu, the non-C-terminal peptides have two carboxy groups at their C-termini, whereas the C-terminal peptides have only one carboxy group. Therefore, using MOLEX, the dicarboxylates of the non-C-terminal peptide form a stable chelate with metal atoms, allowing the isolation of the more weakly retained C-terminal peptides. On the other hand, the CHAMP-NC method consists only of tryptic digestion followed by SCX separation. When digested by trypsin, which cleaves the C-terminal side of Lys and Arg, the non-terminal peptides have more than +2 charge while the acetylated N-terminal and C-terminal peptides have +1 charge under acidic conditions. Therefore, using SCX, the acetylated N- and C-terminal peptides, which are weakly retained, can be isolated because the non-terminal peptides with two positive charges are strongly retained by SCX. Note that, unlike the CHAMP-N method, unmodified N-terminal peptides with +2 charge are not isolated by this method.

**Figure 5.**
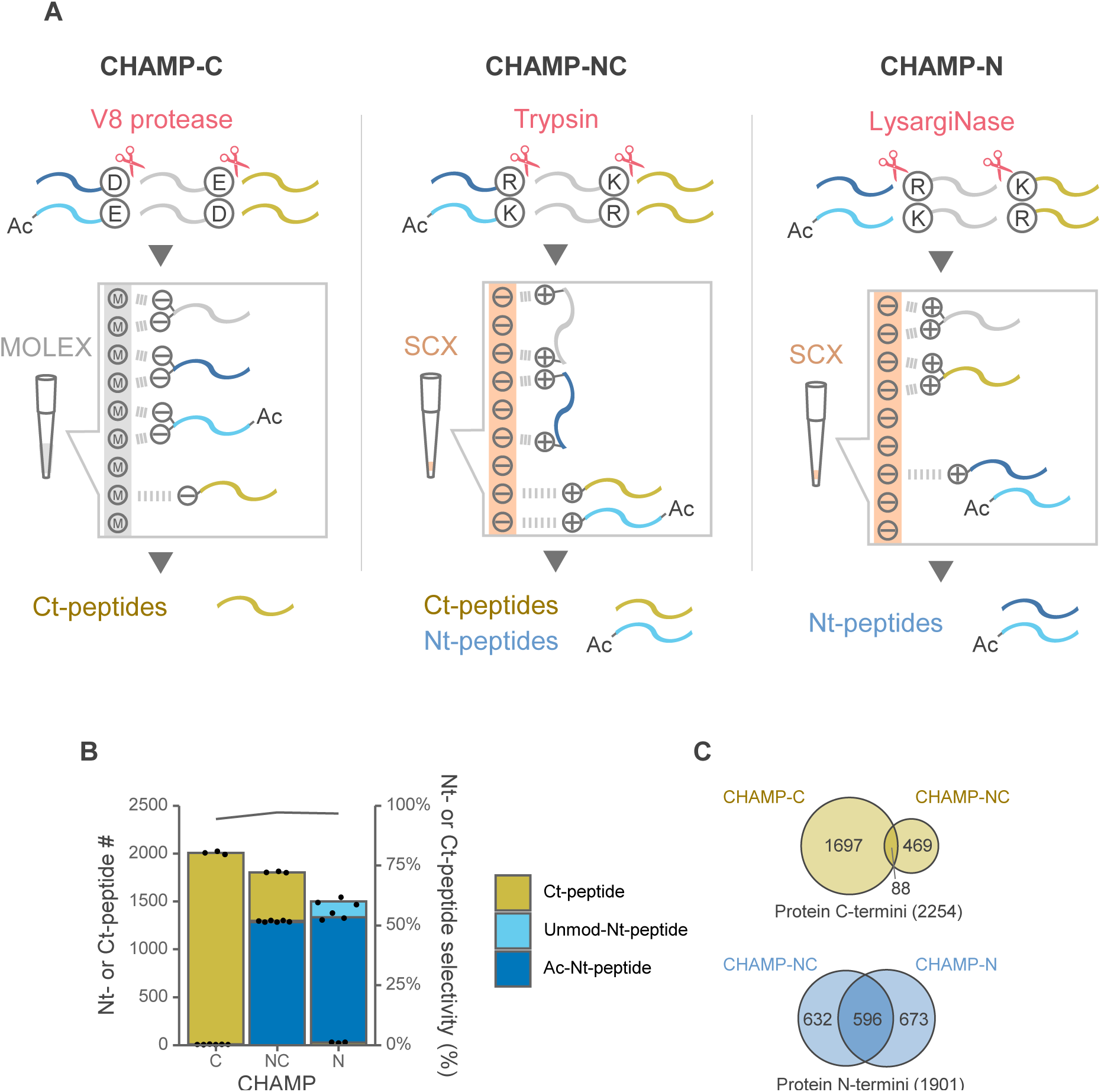
Comprehensive analysis of protein N- and C-termini using the three CHAMP methods. The Orbitrap Exploris 480 system was used. (A) Schematic illustration of protein N- and C-terminal peptide enrichment. (B) Numbers of identified acetylated N-, unmodified N- and C-terminal peptides. N- or C-terminal peptide selectivity (%) was calculated as the sum of the signal intensity of all identified N- and C-terminal peptides divided by that of all identified peptides. (C) Overlaps of protein groups identified with N- or C-terminal peptides in each CHAMP method. Peptides identified in at least one of the three replicates were analyzed.

We performed a comprehensive protein terminome analysis of HEK293T cell-derived proteins using these three CHAMP methods (Figure 5B). In each CHAMP method, HEK293T cell-derived proteins were analyzed in triplicate, and 10 µg of protein sample was used for terminal peptide isolation in each replicate. In total, 2254 protein C-termini and 1901 protein N-termini were identified (Figure 5C). Since each CHAMP method is based on a different digestion enzyme and isolation mechanism, the overlap rate in identified protein termini between methods is low, and the combination of the three CHAMP methods provides a more comprehensive analysis than using each alone (Figure 5C). As for protein C-termini, C-terminal peptides identified by the CHAMP-NC method tend to have acidic amino acid residues near the peptide C-termini, and such C-terminal peptides are not easily identified by the CHAMP-C method because they are relatively strongly retained by MOLEX (Figure S6A). On the other hand, C-terminal peptides with Lys and Arg near the peptide C-termini are too short to be identified by the CHAMP-NC method using trypsin digestion but can be easily isolated by CHAMP-C method (Figure S6B). As for protein N-termini, many N-terminal peptides identified only by the CHAMP-N method are not acetylated or have extra basic amino acids, i.e., extra positive charge (Fig. S6C-E). Trypsin-digested N-terminal peptides always have Lys or Arg, while LysargiNase-digested N-terminal peptides have less positive charge derived from Lys and Arg. Therefore, with the CHAMP-N method, it is possible to isolate N-terminal peptides even with extra positive charge by SCX. For LysargiNase-digested N-terminal peptides identified by the CHAMP-N method, the effect of net charge (Z) in acidic solution on the charge state of the precursor ion in gas phase, the number of b- and y-ions after fragmentation, and the Andromeda score in database searches by MaxQuant was investigated (Fig. S6F-G). As reported previously (13), the charge state of the precursor ion was above +2 even for N-terminal peptides with Z = 0. Also as Z increased, the number of product ions per spectrum and the Andromeda score tended to increase slightly, possibly because the peptides also became longer as Z increased due to missed cleavage. On the other hand, trypsin-digested N-terminal peptides identified by the CHAMP-NC method are always Z ≥ 1. Therefore, combining terminomics methods based on different digestion enzymes and isolation mechanisms can improve coverage and allow for more comprehensive identification of protein termini.

## Conclusions

We have succeeded in developing a tip-based CHAMP-N method that can be parallelized while retaining the high performance (identification number and selectivity) of the HPLC method. We have applied this method to identify novel post-translational cleavages (signal peptides and transit peptides) and non-canonical translation initiation sites, including initiation from near-cognate codons. In combination with the other two tip-based CHAMP methods, one-step terminomics can now be performed very easily. The CHAMP methods are expected to become a widely used standard method for terminomics in the future.

## Data Availability

The raw MS data and analysis files have been deposited with the ProteomeXchange Consortium (http://proteomecentral.proteomexchange.org) via the jPOST partner repository (https://jpostdb.org/) with the data set identifier PXD048917.

## Supplemental data

This article contains supplemental data.

## ACKNOWLEDGEMENTS

We would like to thank the members of the Department of Molecular Systems BioAnalysis for fruitful discussion.

## Funding and additional information

This work has been funded by the JST Strategic Basic Research Program CREST (No. JPMJCR1862), AMED-CREST program (No.JP18gm1010010), and JSPS Grants-in-Aid for Scientific Research 21H02459, 23K18185to Y. I., 20H03241 to K. I. and 23H04924 to Y.I. and K. I., and JST FOREST (No. JPMJFR214L) to K.I.

## Author contributions

K.M., H.N., K.I., and Y. I. designed research; K.M., and H.N. performed research; K.M. analyzed data; K.M., H.N., K.I., and Y. I. wrote the paper.

## Abbreviations

The abbreviations used are:

SCX: strong cation exchange
CHAMP-N: chromatographic amplification of protein N-terminal peptides
ACN: acetonitrile
COFRADIC: combined fractional diagonal chromatography
TAILS: terminal amine isotopic labeling of substrates
CHAMP-C: chromatographic amplification of protein C-terminal peptides
PTS: phase-transfer surfactants
SDC: sodium deoxycholate
SLS: sodium *N*-lauroylsarcosinate
Tris−HCl: tris(hydroxymethyl)aminomethane hydrochloride
TFA: trifluoroacetic acid
AGC: automatic gain control
CV: compensation voltage
FA: formic acid
iMet: initiator methionine
CHAMP-NC: chromatographic amplification of protein N- and C-terminal peptides
MOLEX: metal oxide-based ligand-exchange
Nt-peptide: N-terminal peptide.

---

**Figure S1.**
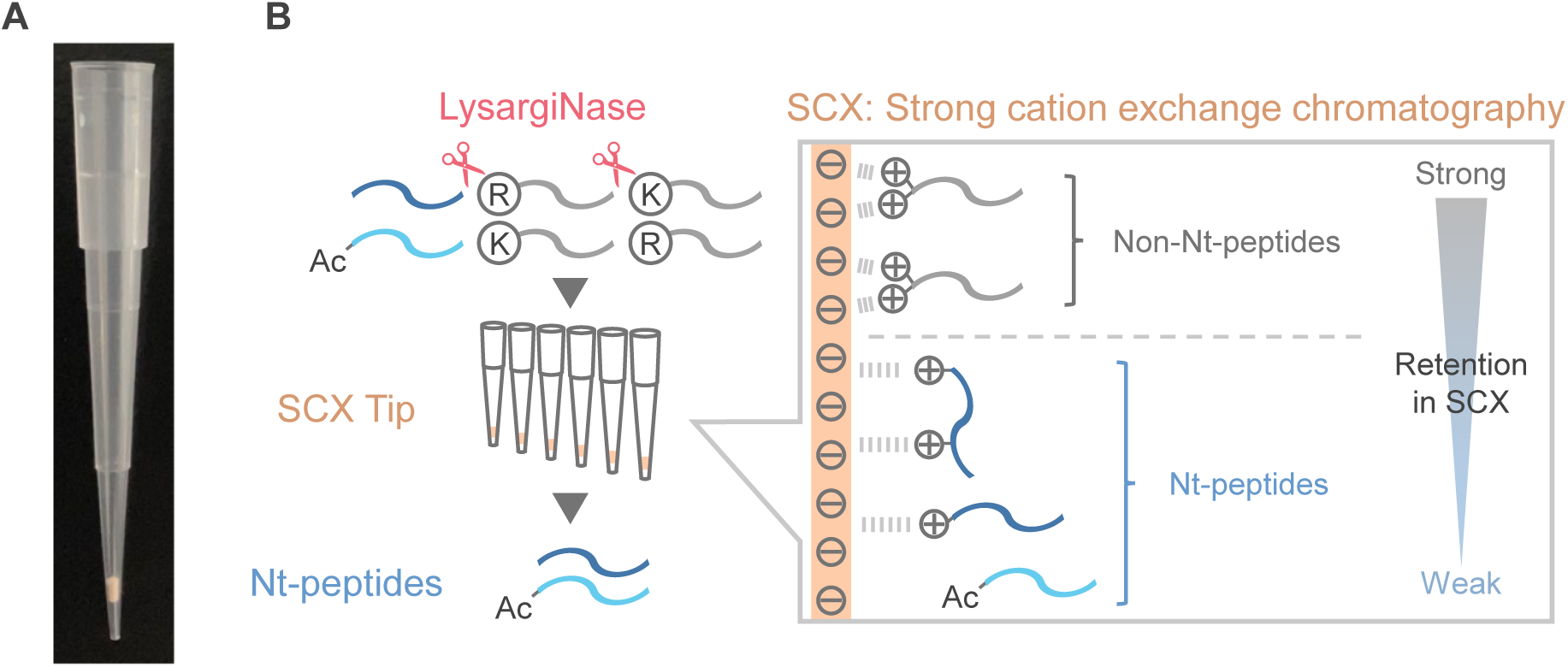
Schematic illustration of protein N-terminal peptide enrichment. (A) Photograph of an SCX-StageTip column. (B) Proteins are digested with LysargiNase, and tip-based SCX chromatography is performed to trap the non-N-terminal peptides. The non-N-terminal peptides are retained in the SCX column, whereas the protein N-terminal peptides flow through the SCX column.

**Figure S2.**
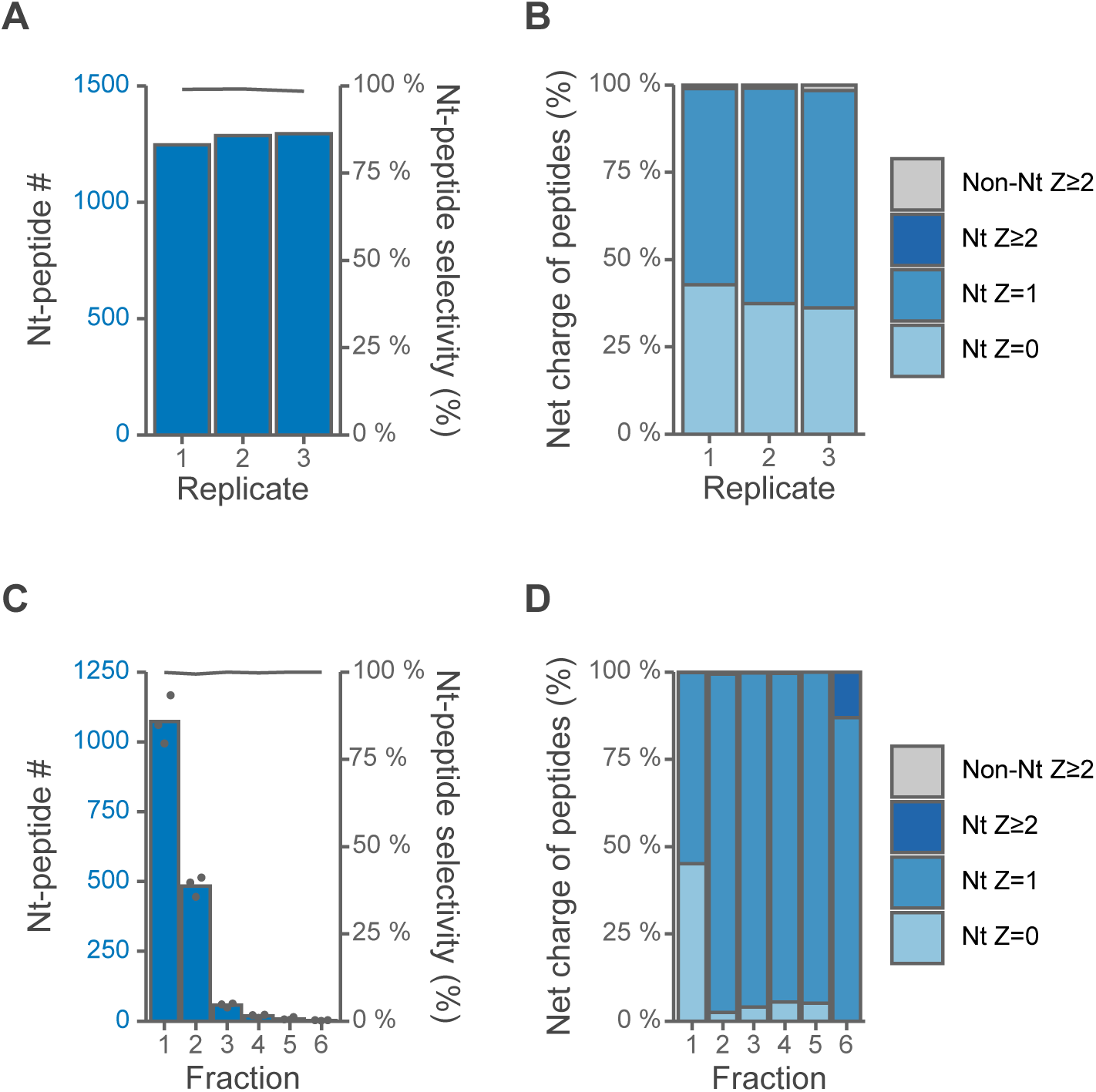
Insufficient separation of protein N-terminal peptides under formic acid-based isocratic condition. SCX tip-based separation of LysargiNase-digested HEK293T peptides under formic acid-based isocratic condition. Z is the charge number at acidic pH, which is based on the number of basic residues per peptide, such as unmodified N-terminus, Lys, Arg and His. The Orbitrap Fusion Lumos system was used. (A) Numbers of identified protein N-terminal peptides. N-Terminal peptide selectivity (%) was calculated as the sum of the signal intensity of all identified N-terminal peptides divided by that of all identified peptides. (B) Distributions of Z values of identified peptides. (C) Numbers of identified protein N-terminal peptides with fractionation.(D) Distributions of Z values of identified peptides with fractionation.

**Figure S3.**
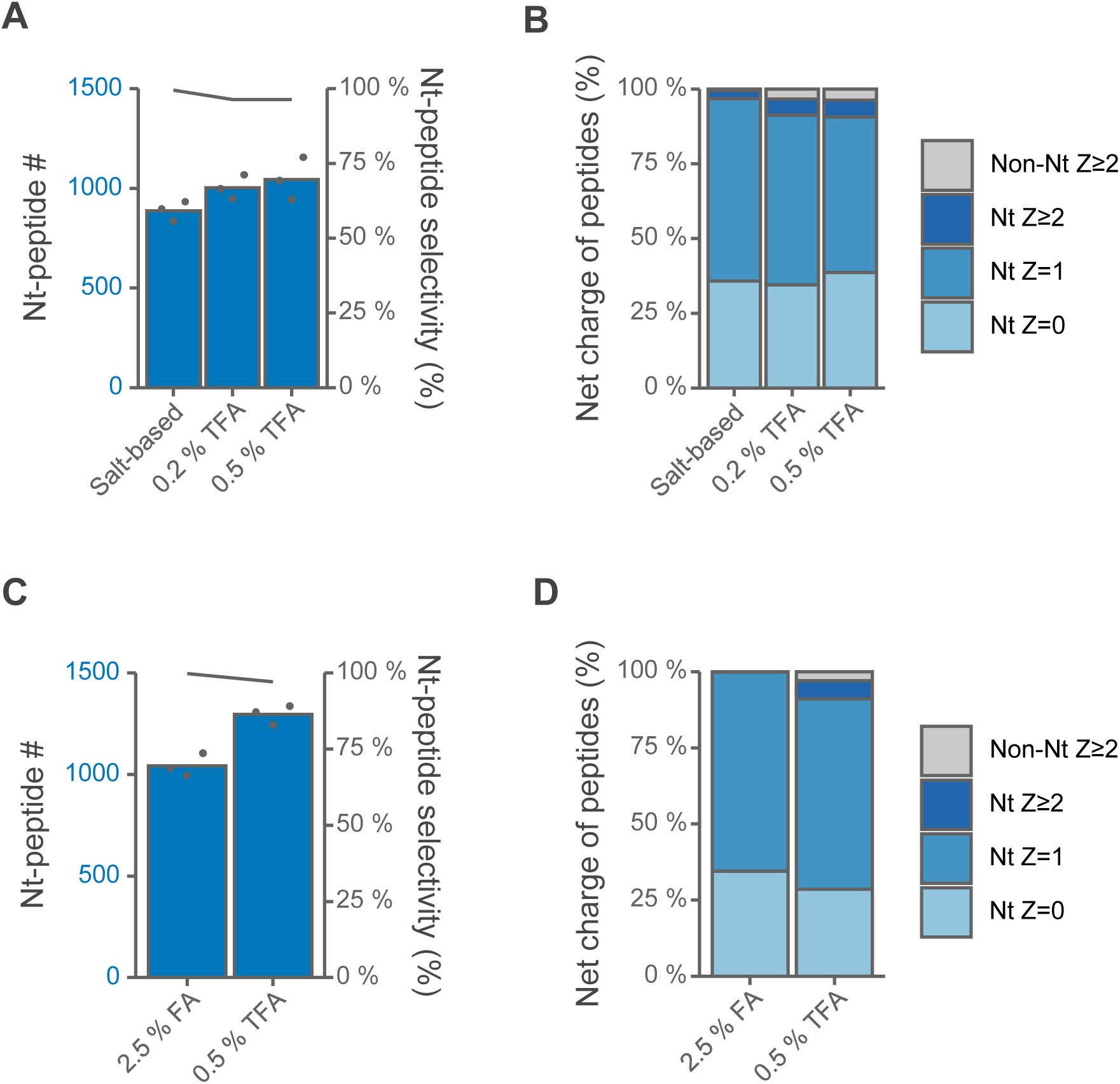
Comparison of salt-based and acid-based isocratic elution of protein N-terminal peptides without fractionation. SCX tip-based separation of LysargiNase-digested HEK293T peptides under isocratic conditions with salt or acid. Z is the charge number at acidic pH, which based on the number of basic residues per peptide, such as unmodified N-terminus, Lys, Arg and His.(A) & (C) Numbers of identified N-terminal peptides. N-Terminal peptide selectivity (%) was calculated as the sum of the signal intensity of all identified N-terminal peptides divided by that of all identified peptides. (B) & (D) Distributions of Z values of identified peptides.

**Figure S4.**
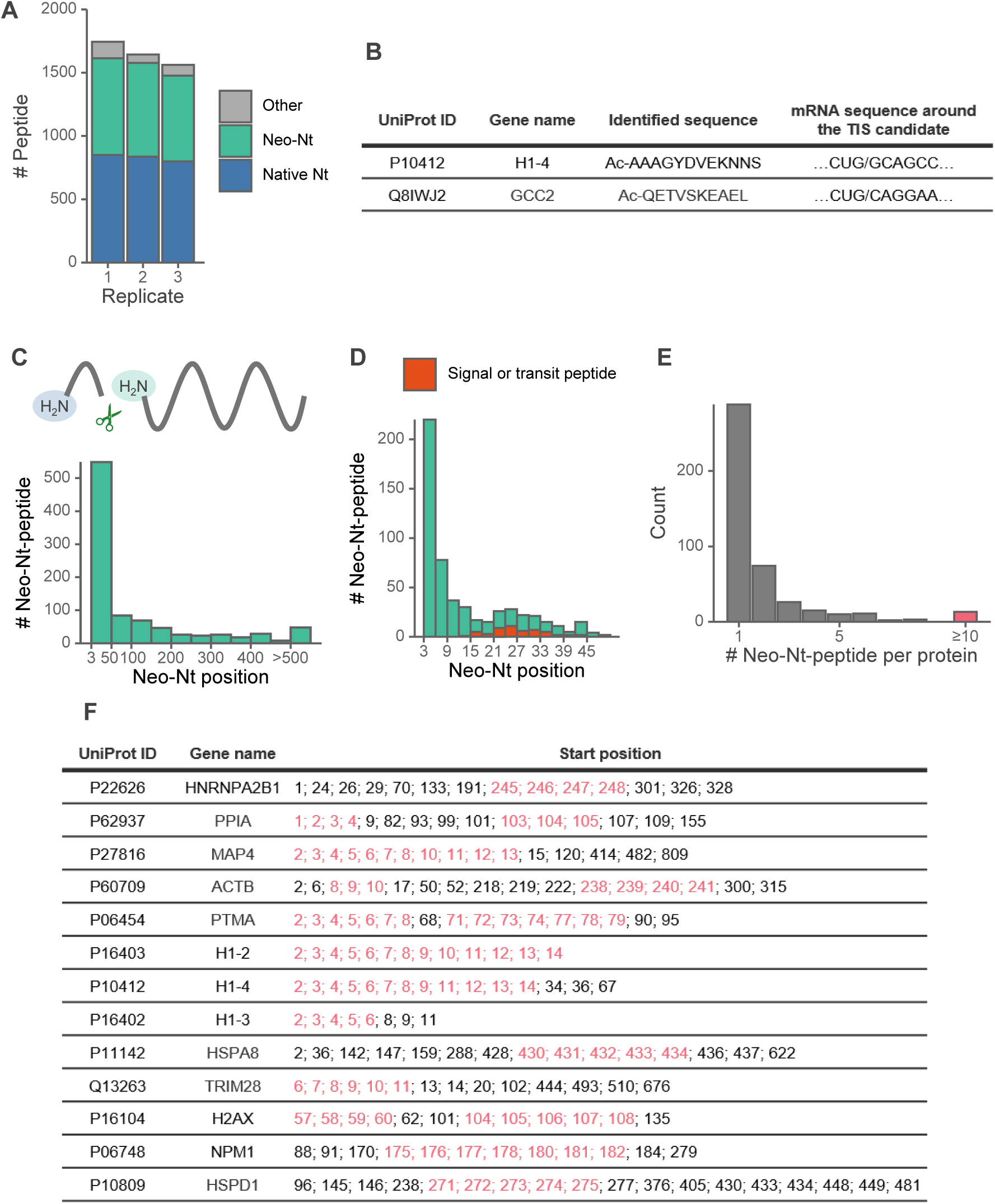
Characteristics of the neo-Nt-peptides. Peptides identified in at least one of the three replicates were analyzed. The Orbitrap Exploris 480 system was used. (A) Numbers of identified peptides with semi-specific search free N-terminus. (B) translation initiation site candidates with near-cognate codons. (C) Neo-Nt starting position of identified neo-Nt-peptides. (D) Zoom into the 3-50 neo-Nt starting position. Transit peptide or signal peptide cleavage site matched peptides are shown in red. (E) Numbers of neo-Nt-peptides per protein. (F) List of proteins with ≥10 neo-Nt-peptides. Peptide ladder sequences (peptides with three or more consecutive start positions) are shown in red.

**Figure S5.**
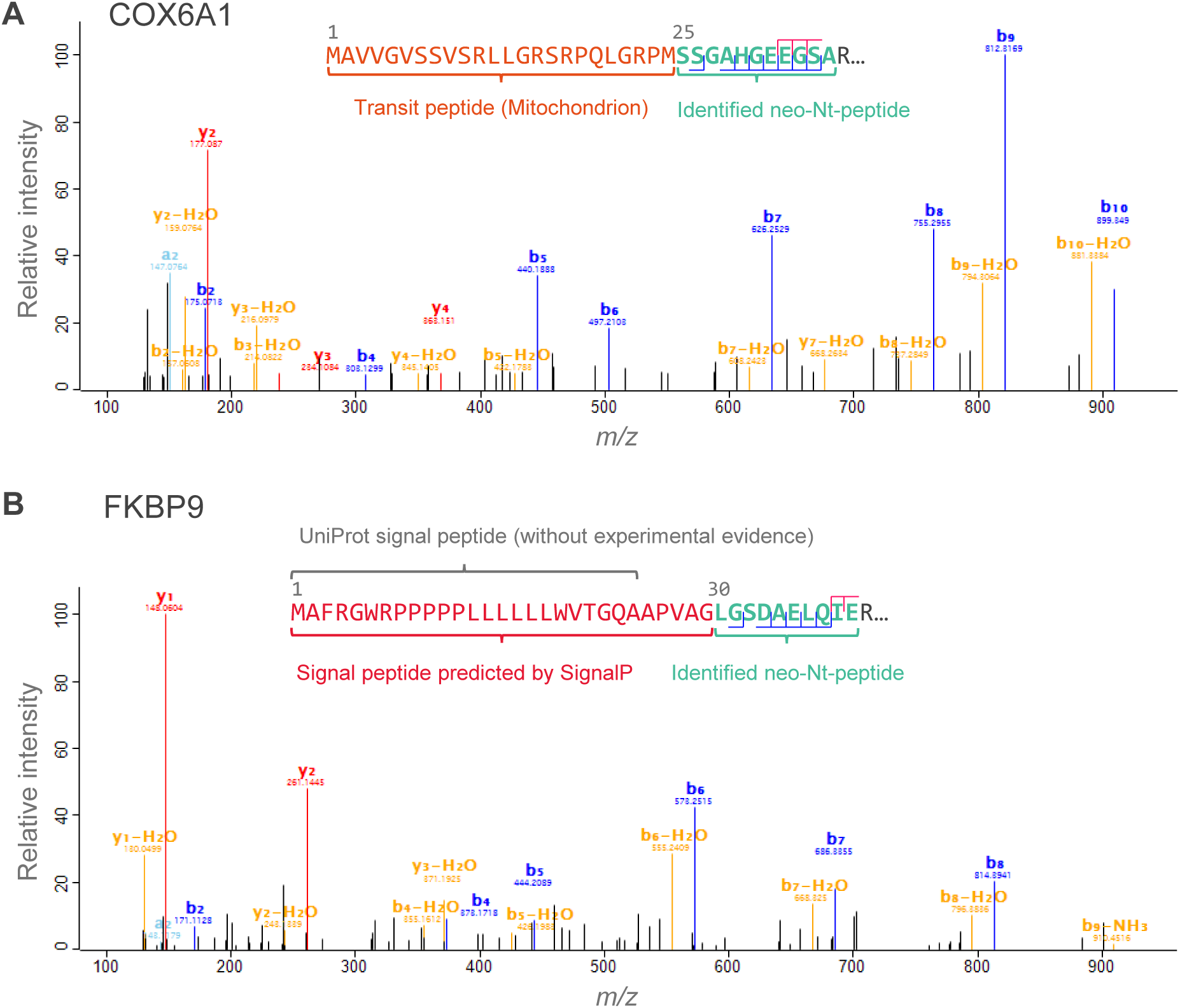
MS/MS spectra of peptides matched to transit peptide or signal peptide cleavage site. The Orbitrap Exploris 480 system was used. (A) MS/MS spectrum of the peptide matched to the transit peptide cleavage site annotated in UniProt. (B) MS/MS spectrum of the peptide matched to the signal peptide cleavage site predicted by SignalP 6.0.

**Figure S6.**
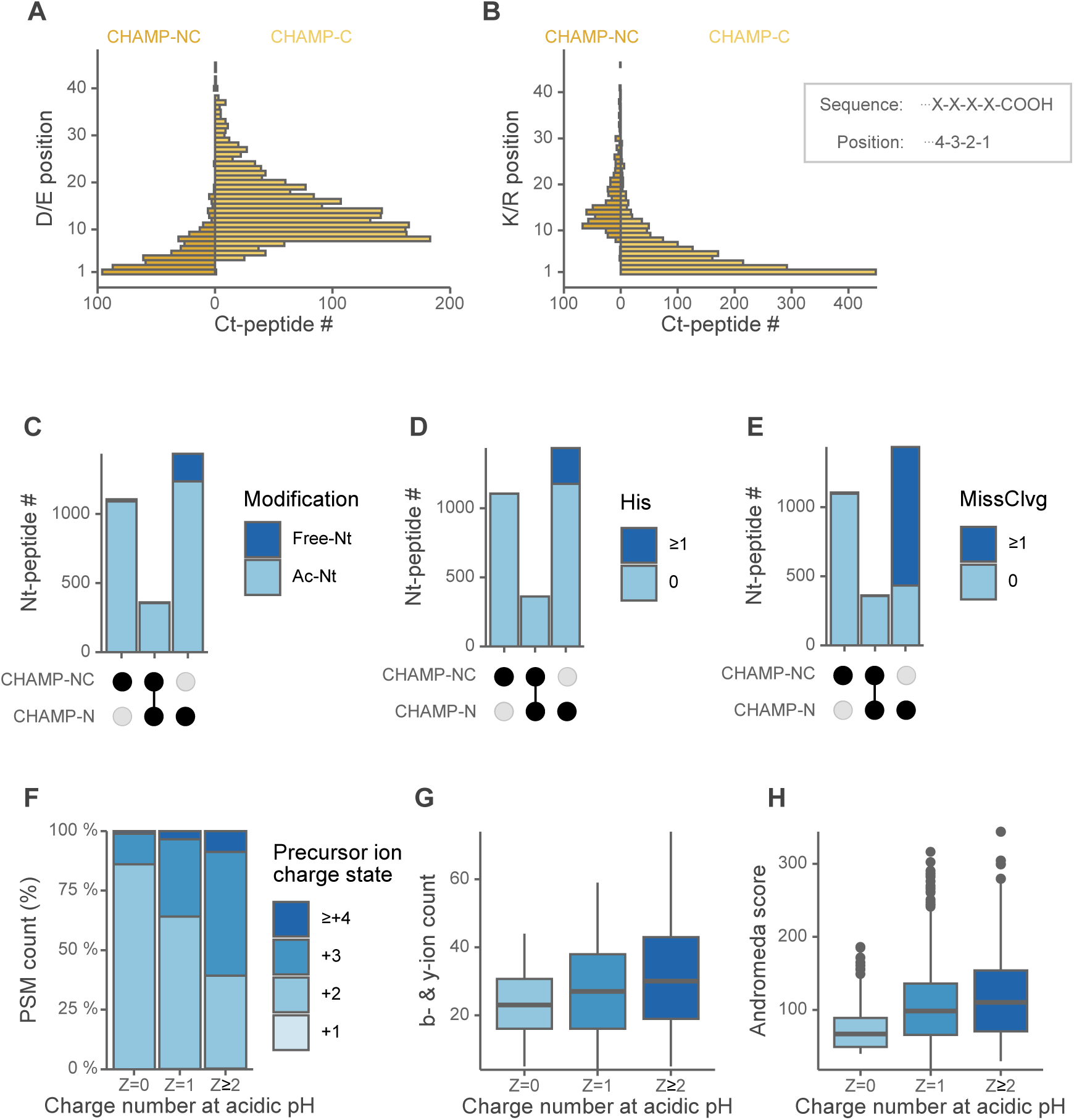
Characteristics of the peptides identified by each of the three CHAMP methods. Peptides identified in at least one of the three replicates were analyzed. The Orbitrap Exploris 480 system was used. (A) The residue distance distribution of the nearest Asp or Glu residue to the identified peptide sequence (including residues adjacent to the V8 protease digestion site) in CHAMP-NC and CHAMP-C experiments. (B) The residue distance distribution of the nearest Lys or Arg residue to the identified peptide sequence (including residues adjacent to the trypsin digestion site) in CHAMP-NC and CHAMP-C experiments. (C) Acetylation states of N-terminal peptides identified by CHAMP-N or CHAMP-NC. (D) Numbers of His residues in the N-terminal peptides identified by CHAMP-N or CHAMP-NC. (E) Number of missed cleavages in the N-terminal peptides identified by CHAMP-N or CHAMP-NC. For peptides identified by both CHAMP-N and CHAMP-NC, missed cleavages in trypsin digested peptides were counted. (F) Distributions of precursor ion charge state of Nt-peptides identified by CHAMP-N. (G) Relationship between charge number at acidic pH and b- and y-ion count of Nt-peptides identified by CHAMP-N. (H) Relationship between charge number at acidic pH and Andromeda score (MaxQuant) of Nt-peptides identified by CHAMP-N.

